# Postnatal environmental enrichment enhances memory by shaping hippocampal-prefrontal theta and gamma rhythms in diploid and trisomic female mice

**DOI:** 10.1101/2022.02.16.480741

**Authors:** Maria Alemany-González, Marta Vilademunt, Thomas Gener, Pau Nebot, M. Victoria Puig

## Abstract

Rich social, physical, and cognitively stimulating lifestyles have powerful effects on cognitive abilities, especially when they are experienced early in life. Cognitive therapies are widely used to attenuate cognitive impairment due to intellectual disability, but also aging and neurodegeneration, however the underlying neural mechanisms are poorly understood. Here we investigated the neural substrates of memory amelioration induced by postnatal environmental enrichment (EE) in diploid female mice and Ts65Dn female mice with partial trisomy of genes ortholog to human chromosome 21, a standard model of Down syndrome (DS, trisomy 21). We recorded neural activities in two brain structures key for cognitive function, the hippocampus and the prefrontal cortex, during rest, sleep and memory performance in mice reared in standard or enriched environments for 7 weeks post-weaning. We found that EE shaped hippocampal-prefrontal neural dynamics in diploid mice and rescued the same disrupted pathways in Ts65Dn mice. The neural activity changes detected in EE-reared wild-type mice combined task-independent adjustments (augmented hippocampal pyramidal activity and gamma synchrony across different brain states) and memory-dependent adjustments (enhanced theta-gamma coupling and ripples in the HPC). Therefore, both brain state adjustments and memory-associated adjustments are good candidates to underlie the beneficial effects of EE on cognition in diploid female mice. Concomitantly, EE attenuated hippocampal and prefrontal hypersynchrony in trisomic females, suggesting distinct neural mechanisms for the generation and rescue of healthy and pathological brain synchrony, respectively, by EE. These results put forward hippocampal hypersynchrony and hippocampal-prefrontal miscommunication as major neural mechanisms underlying the beneficial effects of EE for intellectual disability in DS.

## INTRODUCTION

A stimulating social, physical, and cognitive lifestyle has powerful effects on cognitive abilities and brain function, especially when it is experienced early in life. Cognitive training programs also exert strong influences on cognition and brain activity in healthy adults (Taya et al., 2015) and are the main nonpharmaceutical interventions used to ameliorate cognitive decline due to natural aging or neurodegenerative diseases such as Alzheimer’s disease (Livingston et al., 2017). Moreover, cognitive rehabilitation is the only therapy currently available for people with intellectual disability, including subjects with Down syndrome (DS, trisomy of chromosome 21). In children and adults with DS, cognitive therapy improves executive functions and adaptive functionality (Karaaslan and Mahoney, 2013; Martínez Cué and Dierssen, 2020; de la Torre et al., 2016). Recent findings suggest an adaptive neuroplastic reorganization of DS brains as a result of cognitive training that leads to a more efficient and flexible network activity (Anagnostopoulou et al., 2021). DS brains display increased brain synchrony as assessed by EEG, MEG and fMRI (Anderson et al., 2013; Figueroa-Jimenez et al., 2021; Pujol et al., 2015; Ramírez-Toraño et al., 2021; Velikova et al., 2011), however the exact neural mechanisms involved in the cognitive training-dependent neural activity reorganization have not been elucidated.

Environmental enrichment (EE) mimics a stimulating lifestyle in laboratory animals and has demonstrated beneficial effects on various aspects of brain structure, function, and cognition. It promotes dendritic branching and synaptic plasticity in the hippocampus (HPC) and the prefrontal cortex (PFC)(Hirase and Shinohara, 2014; van Praag et al., 2000), two brain regions crucial for learning and memory, and improves cognitive performance in several tasks such as learning, working memory and recognition memory (Brenes et al., 2016; Leger et al., 2015; Mesa-Gresa et al., 2013; Wang et al., 2020). Studies conducted in anesthetized rodents report that animals reared in enriched environments present enhanced gamma rhythms within hippocampal circuits (Hirase and Shinohara, 2014; van Praag et al., 2000; Shinohara et al., 2013; Tanaka et al., 2017). However, no studies have yet investigated how experience modulates long-range connectivity during wakefulness and natural sleep.

EE has also been found to attenuate molecular, cellular and behavioral deficits in animal models of numerous neurological and psychiatric disorders, including Parkinson’s disease, stroke, depression, schizophrenia and autism spectrum disorders (Nithianantharajah and Hannan, 2006). As expected from the clinical studies, EE also exerts profound changes in the brains and behaviors of mouse models of DS. The Ts65Dn partial trisomic DS model, with triplication of 90 genes ortholog for genes located in human chromosome 21, exhibits behavioral, cellular and molecular phenotypes relevant for DS. Overexpression of genes disrupts neurogenesis and synaptic plasticity in Ts65Dn mice, simplifies dendritic architecture, and imbalances excitation–inhibition that leads to a general overinhibition (Belichenko et al., 2007; Cramer and Galdzicki, 2012; Kleschevnikov et al., 2004; Ruiz-Mejias, 2019). EE decreases inhibition and ameliorates dendritic complexity, synaptic plasticity and cognitive performance in DS mice both during development and adulthood (Begenisic et al., 2011, 2015; Martínez-Cué et al., 2002, 2005; Pons-espinal et al., 2013).

In this study we aimed to unravel the neural substrates of cognitive amelioration induced by EE experienced early in life in diploid healthy mice and trisomic Ts65Dn mice. Ts65Dn mice exhibit poor learning and memory abilities that are greatly improved in females housed postnatally in enriched environments, an effect not observed in Ts65Dn males (Martínez-Cué et al., 2002). Thus, we used female mice in this study. Females were housed in enriched or non-enriched environments for seven weeks after weaning and then neural activity was recorded in the HPC and the PFC during quiet wake, sleep and memory performance. In consonance with the increased brain synchrony reported in clinical studies, we have recently demonstrated hypersynchronous neural activities in hippocampal-prefrontal pathways of Ts65Dn male mice across different brain states that are partially rescued by green tea extracts containing epigallocatechin- 3-gallate (EGCG) (Alemany-González et al., 2020). This allowed us to assess abnormal neural activities in male and female trisomic subjects and compare the actions of EGCG and EE on the same circuits.

## RESULTS

### Female mice reared in enriched environments exhibit increased pyramidal activity and gamma synchrony in the hippocampus during awake resting states

We investigated differential neural dynamics in hippocampal-prefrontal pathways between mice reared in enriched environments (EE group, n = 6 mice) and non-enriched environments (NE group, n = 6 mice) for seven weeks after weaning. Animals reared in enriched conditions were housed in groups of 5 to 7 subjects (mixed diploid and trisomic littermates) in large cages with toys that promote physical activity. The toys were replaced every few days to keep physical and cognitive stimulation constant over weeks. Animals reared in non-enriched conditions were housed in standard cages without toys in groups of 2 or 3 individuals. Following the periods with or without enrichment, animals were implanted with electrodes in the CA1 region of the HPC and the prelimbic medial PFC. After a post-surgical recovery period (typically one week), we recorded neural activities during resting episodes, memory performance in the novel object recognition (NOR) task, and during natural sleep. Recordings during sleep were implemented following the familiarization phase of the memory task to better capture neural signals associated with memory consolidation processes. After the experiments ended, the electrode placements were confirmed histologically (Fig. 1A).

**Figure 1.**
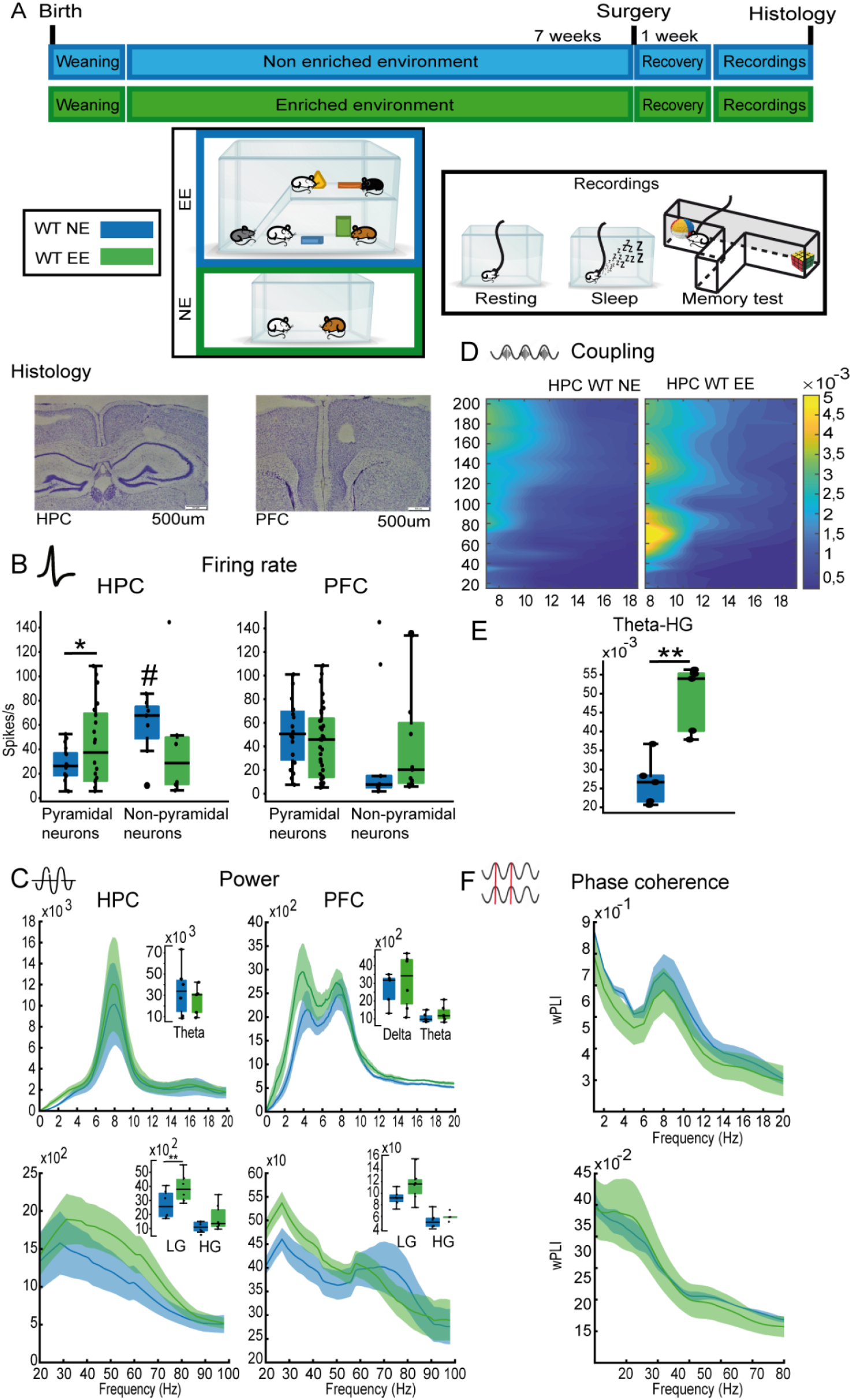
Environmental enrichment enhances pyramidal activity and gamma oscillations in the HPC of diploid healthy females during quiet wakefulness. (A) Experimental timeline. After weaning, mice were reared in enriched or standard environments for 7 weeks. Then, mice were implanted with electrodes in the HPC and PFC and after a recovery period (typically 7 days), behavioral and neurophysiological assessment was conducted during quiet wakefulness, natural sleep and memory performance in the novel object recognition task (NOR). Also shown are representative examples of histological validations of electrodes location in the CA1 region of the HPC and prelimbic PFC. Note the small lesions caused by low-intensity electrical stimulations, which were used to mark the tips of the electrodes after the last recording session in all animals. (B) Mean firing rates of pyramidal and nonpyramidal neurons in HPC and PFC. # indicates significantly different from pyramidal neurons of the NE group. (C) Power spectra of signals in HPC and PFC during quiet wakefulness. (D) Co-modulation maps of cross-frequency coupling in the HPC. Color scale indicates modulation index (MI). (E) Quantification of theta-high gamma MI in the HPC. (F) Phase coherence spectra (wPLI, weighted phase lag index) between the HPC and the PFC of both rearing groups. Data are represented as mean ± SEM. ^*,#^P ≤ 0.05.

We first investigated neural activities during resting states, brief episodes of low behavioral activity that typically lasted from 2 to 4 seconds. We found that the firing rates of individual neurons (SUA) in the HPC and the PFC were similar between rearing groups (mixed ANOVA with brain region and rearing as factors; HPC NE, EE mice; n = 36, 34 neurons, respectively; PFC NE, EE mice: n = 30, 37 neurons, respectively). We further classified neurons as putative pyramidal and non-pyramidal cells based on the shape of their action potentials and their firing pattern (Alemany-González et al., 2020; Kim et al., 2016). Around 70% and 65% of neurons recorded in the HPC and PFC, respectively, were identified as putative pyramidal cells. Presumed pyramidal neurons in the HPC of EE mice fired more action potentials than in NE mice (43.83 ± 7.7 vs. 26.08 ± 3.3 spikes/sec; unpaired T test, P = 0.044), whereas spiking activity in the PFC was similar between groups (41.27 ± 5.9 vs. 46.61 ± 6.8; P = 0.56). Non-pyramidal neurons in the HPC of NE animals displayed higher firing rates than pyramidal neurons, and this increase tended to be reduced in the EE group (Fig. 1B).

Rearing conditions also shaped hippocampal-prefrontal oscillatory dynamics and communication during resting states. Consistent with the augmented activity of hippocampal pyramidal neurons, low gamma power (30-50 Hz) was amplified in the enriched group compared to the non-enriched group in the HPC, but not in the PFC (mixed ANOVA with brain region as within factor and rearing condition as between factor; interaction, F_1,10_ = 9.739, P = 0.011; corrected *post hoc* comparisons, HPC: P = 0.009, PFC: P = 0.11; Fig. 1C). A similar trend was observed for hippocampal high gamma oscillations (50-80 Hz), whereas delta and theta oscillations remained equivalent between groups. To gain deeper insight into the effects of EE on neural synchronization, we quantified the phase-amplitude cross-frequency coupling (PAC) between the phase of slow rhythms (5-20 Hz) and the amplitude of faster oscillations (20-200 Hz) in the HPC (Delgado-Sallent et al., 2021; Jensen and Colgin, 2007; Scheffer-Teixeira and Tort, 2017). We found that in enriched females coupling was strengthened between phases from 5 to 9 cycles/s (theta) and amplitudes from 60 to 80 cycles/s (high gamma; Mann-Whitney test, P = 0.008; Fig. 1D,E). Moreover, circuit synchronization, estimated via phase coherence (weighted phase lag index, wPLI) was not different between rearing groups (Fig. 1F). Together, these results identify relevant roles for hippocampal pyramidal activity and gamma synchrony (power and coupling to theta waves) in the cellular mechanisms of early social, physical, and cognitive stimulation in healthy mice during resting states.

### Environmental enrichment entrains hippocampal gamma oscillations and hippocampal-prefrontal gamma connectivity during memory retrieval

We next investigated how enriched or poor environmental stimulation early in development impacts memory performance and its neural correlates in young adulthood (2- to 3-month-old mice). We used the novel object recognition (NOR) task, a well validated memory test that relies on the mice’ innate instinct to explore novel objects and depends on hippocampal-prefrontal circuits (Alemany-González et al., 2020; Warburton and Brown, 2015). The task was implemented in a T-maze and consisted in a habituation phase, familiarization phase, and memory test phase, each lasting ten minutes. During the habituation phase, the animals explored the maze without objects. Five minutes later, mice were put back in the maze where two identical objects had been placed at the end of the lateral arms for the familiarization phase. Twenty-four hours later, a familiar object and a new object were placed in the maze for the memory test (Fig. 2A). The animals’ interactions with the familiar and the novel objects were temporally aligned to the electrophysiological recordings by pressing two buttons on a joystick for the duration of each visit (Alemany-González et al., 2020). This allowed us to perform a comprehensive characterization of the mice’ behavior and corresponding neural activities in the circuit of interest. We investigated the neural substrates of memory retrieval and novelty/memory acquisition (which cannot be disentangled in this task) by comparing the neural signals recorded during the visits to familiar and novel objects, respectively, during the memory test.

**Figure 2.**
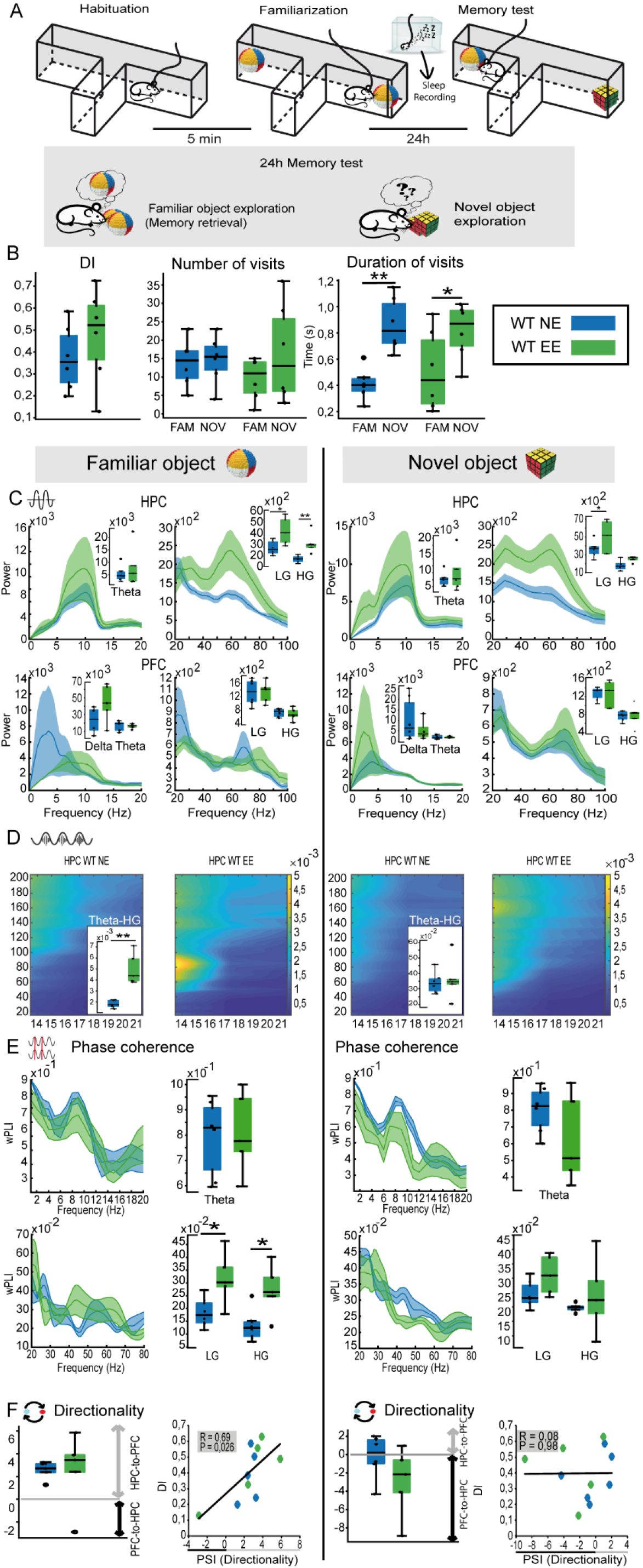
Environmental enrichment enhances hippocampal gamma activity and prefrontal-hippocampal gamma connectivity during memory retrieval in diploid healthy mice. (**A**) The novel object recognition task consists of three phases of 10 minutes each. 1) Habituation phase: mice explore an empty maze. 2) Familiarization phase: mice explore two identical objects placed at the end of the lateral arms of the maze. During this phase, memory for the presented object is first acquired and then stored into memory. 3) Twenty-four-hour memory test: mice explore two objects, one is familiar (presented the previous day) and the other is novel. The arm where the familiar and novel objects are positioned on each session are chosen randomly. During this phase, the memory for the familiar object is retrieved from memory while the memory for the novel object is being acquired. (**B**) Discrimination indices (DIs) and number of visits to familiar and novel objects were similar between rearing groups. In addition, visits to novel objects were longer than to familiar objects in both groups. (**C**) Power spectra of signals in HPC and PFC. Quantifications of relevant bands are also shown. (**D**) Co-modulation maps of cross-frequency coupling in the HPC and corresponding quantification of theta-high gamma MI. (**E**) Circuit’s phase coherence spectra and corresponding quantification at relevant bands. (**F**) HPC-to-PFC directionality (PSI) at gamma frequencies and corresponding correlations with DIs.

Discrimination indices (DIs) for novel versus familiar objects were positive for all the mice ([time visiting the novel object - time visiting the familiar object] / total exploration time; mean DI 0.42 ± 0.05, n = 12 mice); that is, mice explored novel objects more than familiar objects, indicating good recognition memory. As observed in a previous study (De Toma et al., 2019), EE mice tended to display higher DIs than NE mice (0.47 ± 0.09 vs. 0.37 ± 0.06), but this difference did not reach significance (unpaired T test, P = 0.36; Fig. 2B). This was likely due to ceiling effects on behavioral performance. To gain deeper insight into the effects of rearing on memory performance, we quantified the number of visits to each object and the mean duration of individual visits per session. Mice tended to visit novel objects on more occasions than familiar objects during each session (16 vs. 12 visits on average) and the visits lasted longer (840 ± 58 ms for novel objects vs. 460 ± 66 ms for familiar objects; F_1,10_ = 28.95; P < 0.0005) in the two rearing groups (NE mice, P = 0.001; EE mice, P = 0.013; Fig. 2B).

Despite the modest beneficial effects of EE on memory performance, differential neural activities were observed between the two rearing groups. During the visits to familiar objects, EE mice exhibited enhanced low/high gamma power in the HPC (mixed ANOVA, region x rearing interaction: F_1,9_ = 9.105, 14.689; P = 0.015, 0.004; HPC: P = 0.029, 0.007; PFC: P = 0.986, 0.649; Fig. 2C) and strengthening of theta-high gamma coupling (9-14 cycles/s with 70-100 cycles/s; Mann-Whitney, P = 0.008; Fig. 2D). Furthermore, circuit phase coherence at low/high gamma ranges was also more robust in EE mice (P = 0.03; Fig. 2E). During the visits to novel objects, low/high gamma power was also amplified in the HPC of EE mice (F_1,8_ = 5.11; P = 0.05; P = 0.037), however hippocampal theta-gamma coupling and gamma circuit coherence were unchanged (Fig. 2C-E). Together, these observations pointed to an enhancement of gamma coordination within hippocampal microcircuits that likely entrained circuit gamma communication in animals reared in enriched environments, particularly during memory retrieval.

We further investigated any flow of information within the circuit via the phase slope index or PSI (Nolte et al., 2008). We detected a neural information flow at low gamma frequencies that originated in the HPC and traveled to the PFC during the visits to familiar objects, in consonance with our previous observations in diploid male mice. In our previous study, these gamma signals seemed relevant for memory retrieval as they correlated strongly with memory performance (Alemany-González et al., 2020). Here, HPC->PFC low gamma signals recorded during the visits to familiar objects also correlated with the DIs (NE and EE groups combined: Pearson R = 0.69, P = 0.026), confirming that both male and female mice with robust HPC->PFC gamma connectivity exhibit superior memory performance. The strength of this gamma communication was similar between rearing groups (Mann-Whitney, P = 0.42), in accord with their similar DIs (Fig. 2F). Conversely, gamma signals recorded during the visits to novel objects did not show a clear directionality and were not associated with memory performances (Pearson R = 0.008, P = 0.98). In fact, none of the other neurophysiological biomarkers investigated showed any association with memory performances neither during memory acquisition nor retrieval, pointing to a critical role of hippocampus-to-cortex gamma connectivity in recognition memory processes.

### Environmental enrichment enhances hippocampal gamma synchrony and sharp wave ripples during sleep

We recorded neural activities during sleep following the familiarization phase of the NOR task, aiming to capture the processing of recently acquired memories (Fig. 2A). Rapid eye movement (REM) and non-rapid eye movement (NREM) episodes were detected during periods of low behavioral activity (i.e., low accelerometer rates) and eyes closed in an open field as previously reported (Alemany-González et al., 2020). In brief, REM sleep is characterized by prominent theta oscillations in the HPC whereas NREM sleep can be easily identified by the emergence of slow oscillations (<4 Hz) in the cortex and sharp wave-ripple (SWR) events in the HPC (highly synchronous oscillatory activity at 100–250 Hz lasting ∼100-200 ms) (Buzsáki, 2002; Steriade, 2006). Cortical slow waves and hippocampal ripples have been shown to contribute to offline information processing relevant for memory retrieval and consolidation (Buzsáki, 2015; Joo and Frank, 2018) and were therefore investigated here in detail.

During both REM and NREM sleep pyramidal neurons in the HPC of EE mice showed elevated firing rates compared to NE mice (REM: WT NE vs. EE, n = 17, 15 neurons; unpaired T test, P = 0.04; NREM: n = 15, 17 neurons; P = 0.039), similar to wake resting states. In addition, during REM sleep, prefrontal pyramidal neurons also fired more action potentials in EE than NE mice (WT NE vs. EE, n = 16, 18 neurons, P = 0.039). Non-pyramidal neurons displayed higher firing rates than pyramidal neurons in both brain regions, however the spiking activity was not different between rearing groups This was most likely due to the diversity of interneuron subtypes recorded that yielded highly variable firing rates (Fig. 3A).

**Figure 3.**
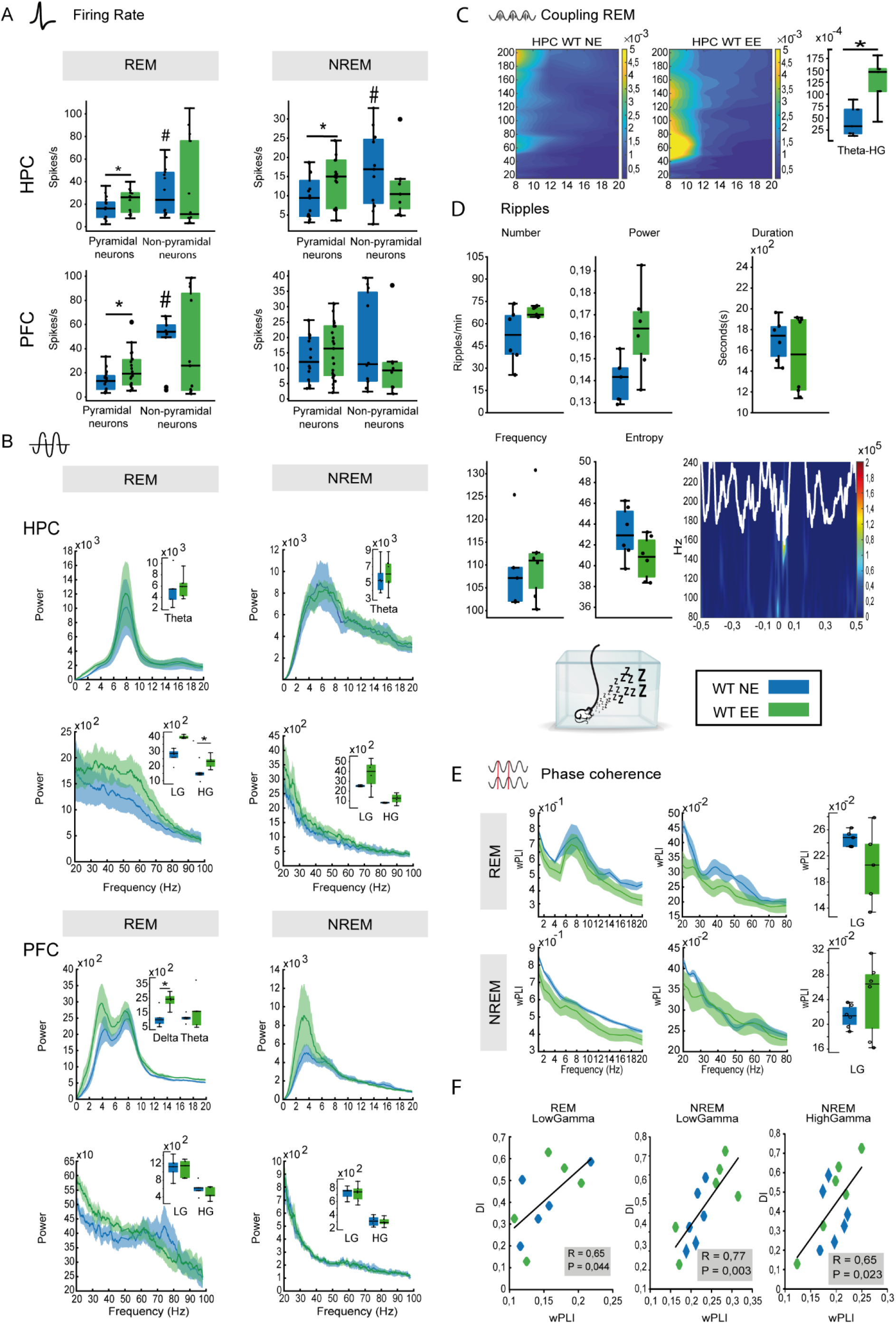
Environmental enrichment increases pyramidal activity and enhances hippocampal gamma synchrony and ripple activity during sleep in diploid healthy mice. (**A**) Mean firing rates of neurons in the HPC and PFC during REM and NREM sleep. (**B**) Power spectra of signals in HPC and PFC during REM and NREM sleep. (**C**) Co-modulation maps of cross- frequency coupling in the HPC during REM sleep and corresponding quantification of theta-high gamma MI. Color scale indicates modulation index (MI). (**D**) Quantification of relevant features of ripple events recorded during NREM sleep. Lower-right panel: representative example of a ripple recorded in a mouse of the WT NE group. (**E**) Circuit’s phase coherence spectra during REM and NREM sleep. Right panels: Quantification of gamma wPLI. (**F**) Correlation between gamma wPLI during REM and NREM sleep and memory performance (DIs).

During REM sleep, and as observed during quiet wake and task performance, the power of gamma oscillations was amplified in the HPC of EE mice (F_1,8_ = 5.959, P = 0.04; Fig. 3B) along with robust strengthening of theta-gamma coupling (8-10 with 40-80 cycles/s; Mann-Whitney, P = 0.01; Fig. 3C). In the PFC, delta power was also enhanced (P = 0.028). During NREM sleep, the power of hippocampal low gamma oscillations and cortical slow waves tended to increase in EE animals, but the differences did not reach significance. To characterize hippocampal ripples, we quantified the number of ripples per minute and their mean power, duration, frequency, and entropy (level of randomness) (Valero et al., 2017). We found that ripple power was slightly larger in EE mice (Mann-Whitney, P = 0.05) while the rest of the parameters were similar between groups (Fig. 3D). Finally, circuit phase coherence was not affected in a consistent way by rearing either during REM or NREM sleep (Fig. 3E). Nevertheless, we found that coherence at high gamma frequencies during REM sleep correlated with DIs (Pearson R = 0.65, P = 0.044; n = 10 mice, NE and EE groups combined) while both low gamma and high gamma coherence during NREM sleep correlated with DIs (R = 0.77, 0.65, P = 0.003, 0.023; Fig. 3F).

### Environmental enrichment prevents pathological theta and gamma hypersynchrony in Ts65Dn female mice during quiet wakefulness

To further understand the pro-cognitive effects of postnatal EE we used a well-established animal model of intellectual disability, the Ts65Dn mouse model of DS. We investigated differential spiking activity, oscillatory power, cross-frequency coupling, and circuit connectivity between wild-type (WT NE: n = 6 mice) and trisomic female mice and further assessed the effects of rearing (TS NE, EE: n = 7 and 6 mice, respectively).

As trisomic males (Alemany-González et al., 2020), trisomic females displayed higher firing rates of pyramidal neurons in the HPC compared to their diploid counterparts during wake resting states (two-WAY ANOVA, genotype x region interaction: F_1,67_ = 13.48, P < 0.0005; WT NE n = 18 vs. TS NE n = 22 neurons; *post hoc* comparison, P = 0.003). This increased hippocampal activity was not observed in the trisomic EE group (TS EE n = 12 neurons vs. TS NE n = 22 neurons, P = 0.002), where firing rates were indeed equivalent to wild-type levels (TS EE vs. WT NE, P = 0.36). The firing rate of pyramidal neurons in the PFC was similar between genotypes and was unaffected by EE (Fig. 4A). Also like TS males, trisomic females exhibited amplified theta and low/high gamma oscillations in the HPC (F_1,21_ = 8.614, 17.273, 11.34, P = 0.008, <0.0005, 0.003; WT NE vs. TS NE mice, P = 0.011, <0.0005, 0.012) as well as in the PFC (F_1,19_ = 7.239, 4.436, P = 0.014, 0.048; WT NE vs. TS NE mice, P = 0.043, 0.014, 0.046; Fig. 4B). We note that excessive theta power in trisomic mice was not due to their inherent hyperlocomotion (Faizi et al., 2011) because during quiet wakefulness the animals did not engage in activities that required mobility. In addition, prefrontal delta waves tended to diminish in TS NE animals, however the changes did not reach significance. Abnormal power attenuated in the TS EE group, where delta, theta and gamma power were close to that observed in diploid animals (TS EE vs. WT NE mice, [HPC theta, gamma] P = 0.151, 0.878; [PFC delta, theta, gamma] P = 0.948, 0.507, 0.168; Fig. 4B). Trisomic females also exhibited broad aberrant cross-frequency coupling between theta waves and oscillations at high frequencies (<80 Hz) in the HPC compared to their non-trisomic peers. Remarkably, theta-gamma coupling was selectively enhanced in TS EE mice in similar profiles than WT EE mice (F_1,18_ = 10.479, P = 0.005; TS EE vs. TS NE, P = 0.005; Fig. 4C), suggesting an improved signal-to-noise ratio of hippocampal synchrony in TS enriched animals. Concomitantly, circuit phase coherence was reduced in trisomic NE mice at high gamma frequencies (F_1,18_ = 16.40, P = 0.001; WT NE vs. TS NE mice: P = 0.005) and partially rescued in the TS EE group (TS EE vs. WT NE mice, P = 0.3; Fig. 4D).

**Figure 4.**
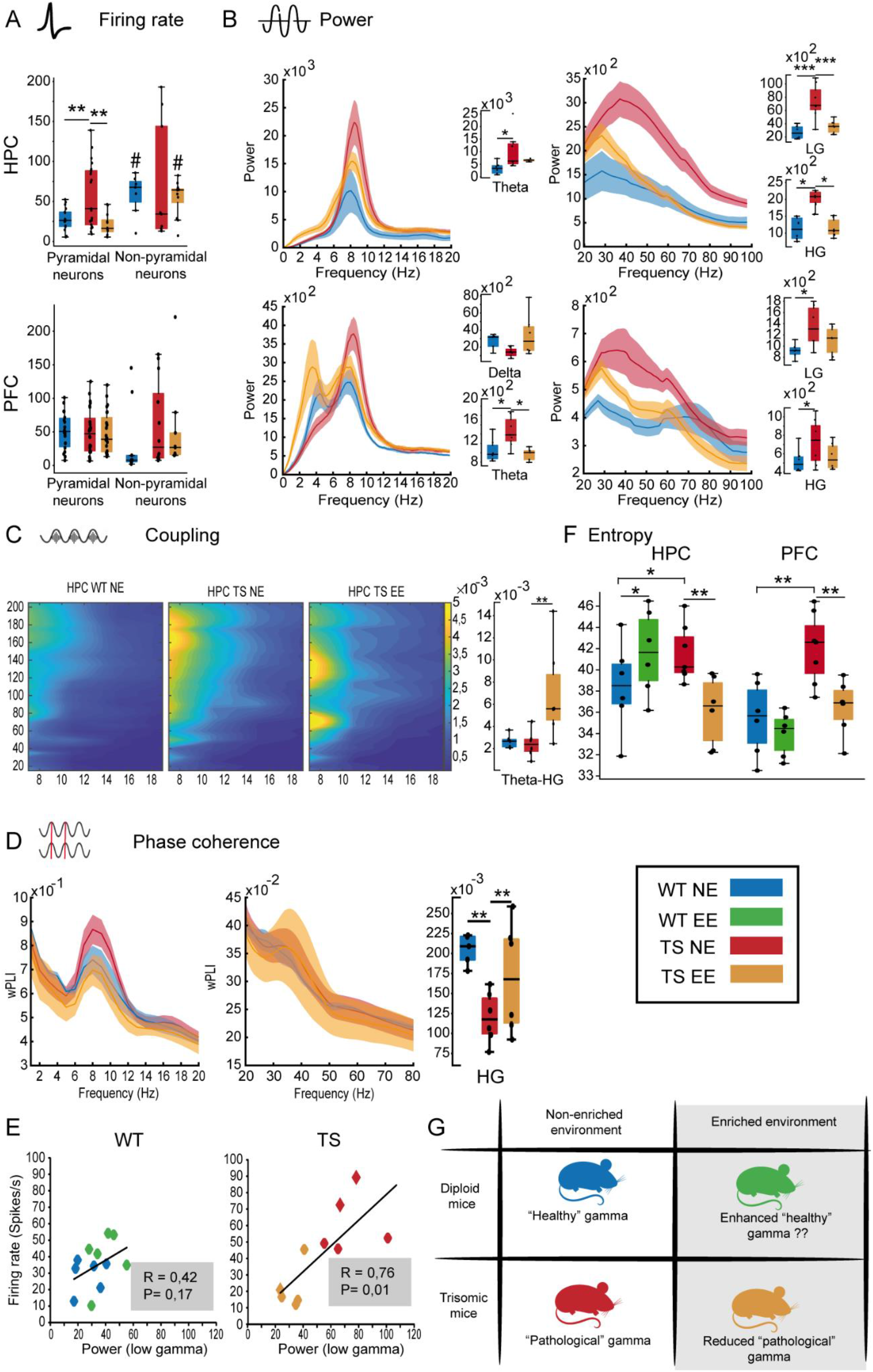
Environmental enrichment prevents pathological theta and gamma hypersynchrony in Ts65Dn female mice during quiet wakefulness. (**A**) Mean firing rates of neurons in the HPC and PFC. The wild-type NE group has been added for reference. (**B**) Power spectra of signals in the HPC and PFC. (**C**) Co-modulation maps of cross-frequency coupling in the HPC and corresponding quantification of theta-high gamma MI. Color scale indicates modulation index (MI). (**D**) Circuit’s phase coherence spectra and corresponding quantification of coherence at high gamma. (**E**) Correlation between firing rate and gamma power recorded in the HPC. (**F**) Quantification of gamma entropy in the HPC and PFC for the four experimental groups. (**G**) Proposed model for the effects of EE on gamma synchrony in diploid and trisomic mice reared in enriched and standard environments.

Taken together, our results indicated opposite effects of EE on gamma rhythms in diploid and trisomic animals. That is, enhancement of hippocampal gamma in enriched wild-types and reduction of excessive gamma in the HPC and PFC of trisomic mice. Based on recent literature (Guyon et al., 2021), we hypothesized that EE favored “healthy” gamma rhythms in diploid animals, gamma synchrony emerging from the interplay between fast-spiking interneurons and pyramidal neurons and reduced “pathological” gamma rhythms in trisomic animals, gamma synchrony generated by the increase of asynchronous or “noisy” firing of excitatory pyramidal neurons due to inhibitory dysfunction. To investigate this, we first looked at whether the spiking activity of pyramidal neurons correlated with gamma power, as it has been proposed that asynchronous firing of neurons can “contaminate” the LFP amplifying gamma oscillations.

In consonance with this, we found that the increased firing rate of pyramidal neurons in the HPC correlated with low gamma power in trisomic mice but not in wild-type mice (Pearson’s correlation; WT NE/EE combined, R = 0.42, P = 0.17; TS NE/EE combined, R = 0.76, P = 0.01; Fig. 4E). We next quantified spectral dispersion by estimating the entropy of the LFP signals. Previous studies have suggested that increased entropy of oscillatory activities may be linked with pathological synchronization in epilepsy and schizophrenia (Guyon et al., 2021; Valero et al., 2017). Trisomic mice displayed increased entropy of low gamma rhythms in the HPC and PFC compared to their diploid littermates (F_1,19_ = 14.76, 15.89, P = 0.001; WT NE vs. TS NE mice: P = 0.027, 0.001) that was reduced in the TS enriched group (TS EE vs. WT NE mice, P = 0.75, 0.58). Interestingly, EE also reduced gamma entropy in the HPC of diploid mice (WT NE vs WT EE, P=0,029; Fig. 4F). Altogether, these results suggested that postnatal EE enhanced “healthy” gamma rhythms in wild-type mice and prevented the emergence of “pathological” gamma rhythms in trisomic animals (Fig. 4G).

### Environmental enrichment reduces excessive theta and gamma synchrony and restores HPC->PFC low gamma connectivity in Ts65Dn female mice during memory retrieval

Here, as previously described, trisomic females exhibited poor memory performances (DIs TS NE vs. WT NE mice, -0.27 ± 0.06 vs. 0.35 ± 0.06; unpaired T test; P <0.0005) and trisomic EE females displayed memory performances similar to the wild-types (TS EE vs. WT NE mice, P = 0.78; Fig. 5A). The negative memory indices in trisomic non-enriched mice resulted from: 1/ the fact that the animals visited the familiar objects more times than the novel objects (16 vs. 13 visits on average per session, respectively), 2/ the visits to novel items were shorter than in diploid mice (509 ms in TS NE mice vs. 840 ms in WT NE mice; P = 0.002) while 3/ the duration of visits to familiar items remained similar (414 ms in TS NE mice vs. 461 ms in WT NE mice; P = 0.23). Together, these behavioral measures likely reflected poor attention of trisomic mice. Notably, the visits of trisomic EE mice to novel objects were longer than to familiar objects (1310 ms vs. 677 ms, respectively; paired T test, P = 0.0005) while they visited novel and familiar objects evenly (P = 0.74; Fig. 5A). The very long duration of visits to novel items by trisomic EE mice (1310 ± 0.17 ms) suggested a strong effect of postnatal EE on attentional abilities.

**Figure 5.**
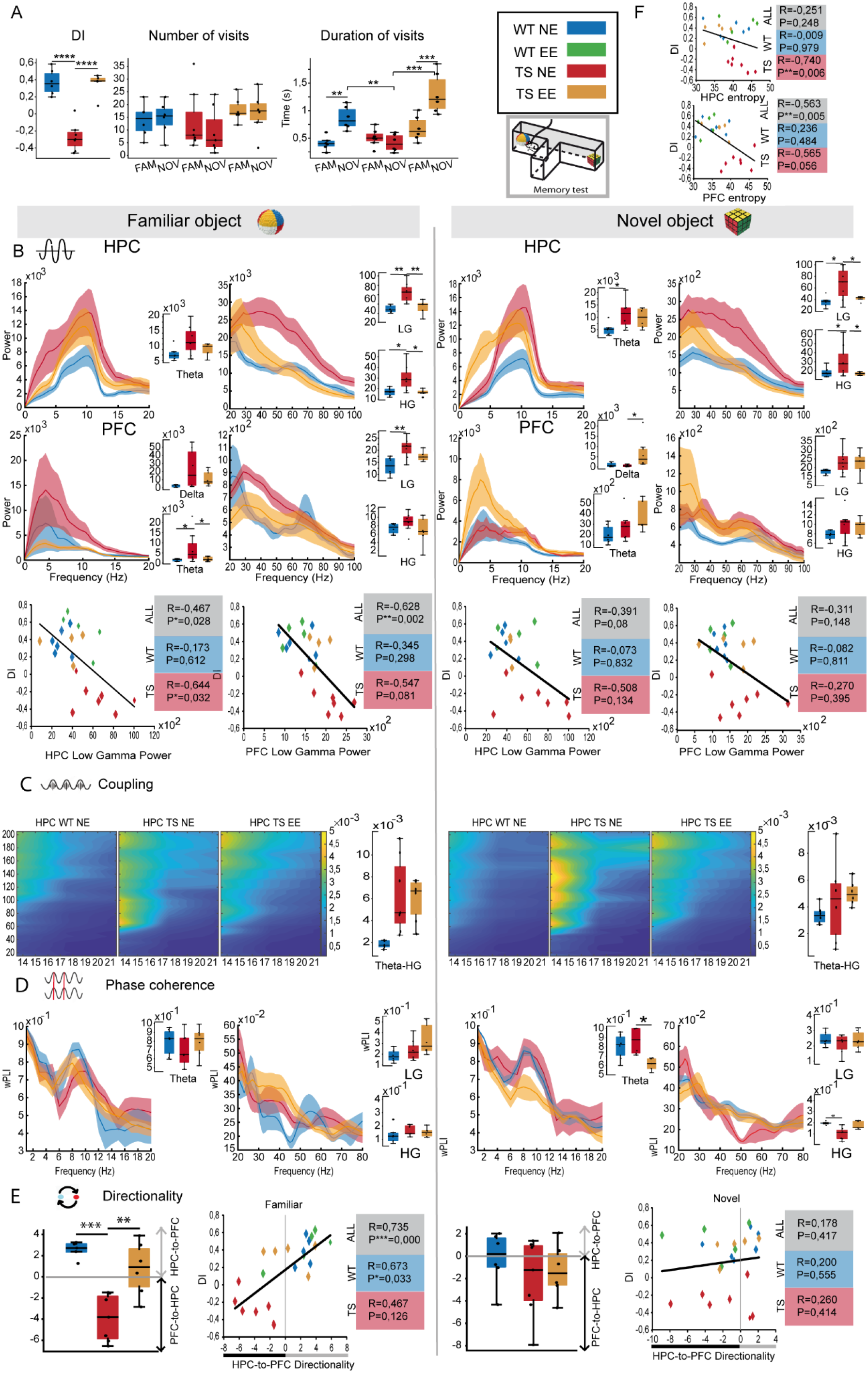
Environmental enrichment reduces excessive theta and gamma synchrony and restores HPC->PFC low gamma connectivity in Ts65Dn female mice during memory retrieval. (**A**) Ts65Dn females exhibited very poor memory performances in the NOR task that were fully rescued in TS enriched animals. This was due to an increase in the duration of visits to objects, particularly the novel items. (**B**) Power spectra of signals in the HPC and PFC during the visits to objects in the maze. Power during visits to familiar and novel objects are depicted on the left and right columns, respectively. Correlations between gamma power in the HPC and PFC and memory performances (DIs) are shown below. (**C**) Co-modulation maps of cross-frequency coupling in the HPC during familiar and novel object explorations. Quantification of theta-high gamma MIs in HPC during familiar and novel object explorations. (**D**) Circuit’s phase coherence spectra during object explorations. Quantifications of relevant bands are also shown. (**E**) HPC-to-PFC directionality (PSI) at low gamma frequencies during object explorations. Corresponding correlations between gamma PSI and memory performances (DIs). (**F**) Correlations between gamma entropy in the HPC and PFC and memory performances (DIs).

We next investigated neurophysiological divergences in Ts65Dn mice during memory retrieval and novelty/memory acquisition. During the visits to familiar objects, trisomic females exhibited increased theta power in the HPC (two-WAY ANOVA, genotype x rearing factors: F_1,19_ = 5.39, P = 0.03) along with amplified low/high gamma power compared with their wild-type peers (F_1,19_ = 13.41, 16.1, P = 0.002, 0.001; TS NE vs. WT NE: P = 0.001, 0.018). As above, theta power increases were not simply due to hyperlocomotion of trisomic mice, as the animals’ mobility was low during the interactions with the objects. The excessive theta and gamma power attenuated in the TS EE group. In the PFC, TS mice showed a less pronounced increase of delta, theta and low gamma power (F_1,19_ = 15.028, P = 0.001) that were also reduced in the EE group (Fig. 5B). Moreover, and similar to resting states, TS NE mice exhibited broad hippocampal coupling between theta and high frequencies (>60 Hz), however theta-high gamma coupling was not enhanced in the TS EE group (F_1,18_ = 10.47, P = 0.005; Fig. 5C). Moreover, theta and gamma phase coherence was similar between diploid NE and trisomic animals reared in both environments (wPLI TS NE vs. TS EE: P = 0.1, 0.7; Fig. 5D). Intriguingly, the selective HPC->PFC flow of information at low gamma frequencies observed in diploid females during memory retrieval reversed in trisomic females, originating in the PFC and traveling to the HPC (F_1,19_ = 16.78, P = 0.001, TS NE vs. WT NE: P = 0.0005). This was attenuated in many trisomic EE mice (F_1,19_ = 5.375, P = 0.032; TS NE vs. TS EE, P = 0.002; Fig. 5E). Collectively, trisomic females housed in standard cages showed aberrant theta and gamma synchrony in both brain areas and abnormal circuit communication at gamma frequencies during memory retrieval that were partially rescued in trisomic animals housed in enriched cages.

During the visits to novel objects, trisomic females also exhibited amplified theta and low/high gamma power in the HPC (F_1,20_ = 5.51, 5.56, P = 0.03; TS NE vs. WT NE: P = 0.01, 0.05). The excessive low/high gamma power was reduced in trisomic EE mice (TS NE vs. TS EE: P = 0.048, 0.038) to wild-type levels (TS EE vs. WT NE: P = 0.36, 0.57), but not the increased theta. In the PFC, theta and low gamma power were again amplified in trisomic NE mice (F_1,20_ = 7.43, 5.44, P = 0.013, 0.03), a tendency that was maintained in EE animals (Fig. 5B). Overall, we found a consistent increase of theta and gamma power in trisomic mice during quiet wakefulness (Fig. 4B) and the NOR task that indicated a pathological hypersynchronization within the HPC and the PFC across brain states. Moreover, hippocampal cross-frequency coupling was again broadly enhanced in trisomic NE animals between theta and high frequencies and this non-specific hypersynchronization diminished in the EE group (Fig. 5C). Phase coherence was weaker at high gamma frequencies in non-enriched TS mice (F_1,19_ = 7.31, P = 0.014; TS NE vs. WT NE: P = 0.023), like in resting states, and this was partially rescued in EE animals (TS NE vs. TS EE: P = 0.07; TS NE vs. WT NE: P = 0.29; Fig. 5D). Finally, we did not observe any difference in the directionality of signals within the circuit between genotypes nor an effect of rearing (Fig. 5E).

As above, we investigated neurophysiological biomarkers relevant for memory performance. Several measures recorded during the visits to familiar objects, but not to novel objects, correlated with the memory indices when combining all animals (n = 24 mice). First, the power of low gamma oscillations in the HPC correlated inversely with the DIs ([familiar, novel visits] R = -0.47, -0.39, P = 0.028, 0.08). This correlation remained significant for the trisomic group (n = 13 mice) ([familiar, novel visits], R=-0.64, -0.51, P=0.032, 0.13) but not for the diploid group (n = 11 mice). Similar associations were observed in the PFC ([familiar, novel visits], R = -0.63, -0.31, P = 0.002, 0.15). This suggests a task-dependent pathological gamma hypersynchrony during the exploration of familiar objects in trisomic mice with potential contribution to their memory impairment (Fig. 5B). Furthermore, and again as previously reported for male mice, HPC->PFC low gamma signals correlated strongly with DIs during visits to familiar objects when combining all the animals (R = 0.74, P < 0.0005; Fig. 5E), unraveling a negative impact of disordered HPC-PFC signaling for memory retrieval in trisomic animals. This correlation remained significant for the diploid group but not for the trisomic group (R = 0.67, 0.47, P = 0.033, 0.13, respectively). In summary, we could establish strong associations between pathological gamma rhythms and poor memory retrieval in trisomic animals that were not found in trisomic mice reared in enriched environments. This was further supported by significant correlations between gamma entropy in the HPC and PFC of TS NE animals and their poor memory performances (R = -0.74, P = 0.006; R = -0.57, P = 0.056; Fig. 5G).

### Environmental enrichment rescues enhanced hippocampal-prefrontal gamma synchrony and ripple activity in Ts65Dn mice during sleep

During episodes of REM sleep, the firing rate of pyramidal neurons was increased both in the HPC and the PFC in trisomic mice compared to their wild-type peers (two-WAY ANOVA, F_1,67_ = 6.05, 5.74, P = 0.017, 0.019; *post hoc* comparisons, P = 0.008, <0.0005). This elevated activity was reduced by EE only in the PFC (P = 0.042; Fig. 6A). Consistent with wake states, trisomic mice also showed augmented delta, theta and gamma oscillations in the HPC and PFC. Interestingly, gamma, but not theta nor delta, oscillations were partially prevented in enriched animals, particularly in the HPC (Fig. 6B). Hippocampal coupling and circuit’s phase coherence remained similar between genotypes and were not influenced by the type of rearing (Fig. 6C,D).

**Figure 6.**
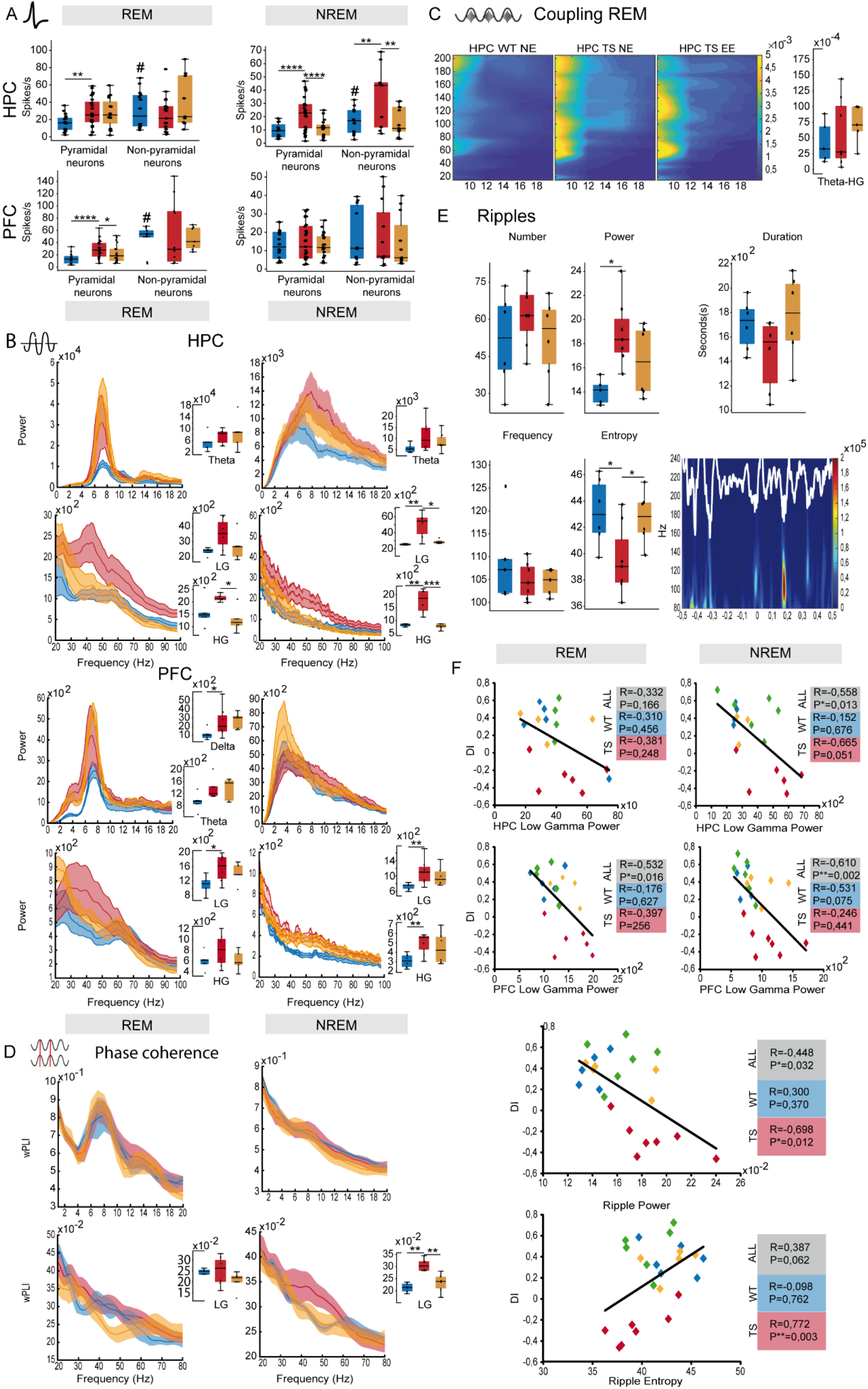
Environmental enrichment rescues enhanced hippocampal-prefrontal gamma synchrony and ripple activity in Ts65Dn mice during sleep. (**A**) Mean firing rates of neurons in the HPC and PFC during REM and NREM sleep. The wild-type NE group has been added for reference. (**B**) Power spectra of signals in the HPC and PFC with quantification of relevant bands. (**C**) Co- modulation maps of cross-frequency coupling in the HPC and corresponding quantification of theta-high gamma MI. Color scale indicates modulation index (MI). (**D**) Circuit’s phase coherence spectra and corresponding quantification of coherence at high gamma. (**E**) Quantification of relevant features of ripple events recorded during NREM sleep. Lower-right panel: representative example of a ripple recorded in a mouse of the TS NE group (same scale as the color plot shown in Fig. 3D). (**F**) Correlations between low gamma power and memory performances (DIs) in both brain regions including the four experimental groups. Similar correlations were detected for high gamma power (not shown). Also shown are correlations between ripple power and entropy and DIs.

During NREM sleep, several neurophysiological anomalies were also detected in trisomic NE mice. First, the firing rates of both pyramidal and non-pyramidal neurons in the HPC were increased (F_1,36_ = 5.39, 5.59, P = 0.023, 0.024; *post hoc* comparisons, P < 0.0005, 0.008). This hippocampal hyperactivity was not observed in EE mice, where firing rates were similar to wild-type levels (TS EE vs. WT NE: pyramidal and non-pyramidal activity, P = 0.38, 0.85; Fig. 6A). In addition, and again, low/high gamma activities in the HPC were increased in trisomic NE animals (F_1,16_ = 8.07, 17.52, P = 0.012, 0.001; P = 0.008, 0.001) and, as observed in other brain states, were fully recovered in enriched mice (P = 0.071, 0.45; Fig. 6B). In the PFC, slow waves were not different between any of the groups. Low/high gamma oscillations were amplified in trisomic mice (F_1,21_ = 11.73, 13.45, P = 0.003, 0.001; P = 0.009, 0.008), however this enhancement was still detected in trisomic EE mice (Fig. 6B). Furthermore, ripple events exhibited enlarged power and reduced entropy in trisomic mice (F_1,16_ = 4.87, 9.23, P = 0.042, 0.008) that were normalized in EE animals (P = 0.096, 0.75; Fig. 6E). Together, these observations point to a main effect of EE on hippocampal microcircuits in trisomic animals during NREM sleep. Moreover, circuit phase coherence was strengthened in trisomic mice at low gamma frequencies (F_1,20_ = 10.1, P = 0.005; P = 0.001) and normalized in trisomic EE mice to wild-type levels (P = 0.29; Fig. 6D). Thus, again, we observed increased gamma synchrony within the circuit during both sleep states that were greatly reduced in trisomic animals reared in enriched environments.

Remarkably, some key neurophysiological biomarkers recorded during sleep were able to predict the animals’ memory performance. Low/high gamma power showed negative associations with DIs both in the HPC and the PFC when combining all groups. The correlations were strongest for prefrontal gamma power during REM sleep ([low/high gamma] R = -0,532, -0,490, P = 0.016, 0,024, respectively) and for prefrontal gamma power during NREM sleep (R = -0.610, -0.550, P = 0.002, 0.005). High Gamma power during NREM sleep also correlated negatively with DI (R = -0.541, P = 0.014). This correlation remained significant in the trisomic group when considered alone, but not in the wild-type group, underscoring even further their pathological nature. Notably, during NREM sleep, ripple power and entropy also correlated with the DIs in the trisomic group (R = -0.698, 0.772, P = 0.012, 0.03; Fig. 6F). Altogether, we unravel pathological spiking, gamma synchrony and ripple events in hippocampal-prefrontal circuits of trisomic mice during sleep that are strongly associated with poor memory and rescued in animals reared in enriched environments.

## DISCUSSION

We report that physical, social, and cognitive stimulation during critical developmental periods shapes hippocampal-prefrontal neural dynamics in wild-type mice and prevents the disruption of the same pathways in Ts65Dn mice. Diploid animals reared in enriched environments exhibited enhanced gamma rhythms in the HPC. Conversely, trisomic mice displayed excessive theta and gamma rhythms in the HPC and PFC that were reduced by EE. We hypothesize that distinct neural mechanisms may be responsible for enrichment-dependent enhancement of “healthy” gamma rhythms in diploid animals and enrichment-dependent reduction of “pathological” gamma rhythms in trisomic animals.

Wild-type mice reared in stimulating environments showed augmented neural activity and gamma synchrony in hippocampal microcircuits across different brain states. More specifically, elevated spiking activity of pyramidal neurons was detected during wake resting states and REM/NREM sleep states. We note that cells classified here as putative non-pyramidal neurons likely included a variety of interneuron subtypes, for which it was difficult to obtain consistent effects. Concomitantly, gamma synchrony (oscillatory power and coupling with theta waves) was enhanced in the HPC consistently during resting states, memory performance, and REM sleep. These results are in line with previous observations reporting amplified gamma oscillations in the HPC of anesthetized rodents reared in enriched environments (Shinohara et al., 2013; Tanaka et al., 2017) and provide new evidence for experience-dependent enhancement of hippocampal gamma synchrony during alertness, sleep, and memory performance. Furthermore, our observations of enhanced ripples during slow wave sleep in the EE group were also reported previously in anesthetized mice, where an implication of astrocytes was demonstrated (Tanaka et al., 2017). Here, ripples were recorded immediately after the familiarization phase of the memory task, thus it is plausible that their enhancement reflected more efficacious processing of events associated with memory acquisition and consolidation in enriched animals (Buzsáki, 2015; Joo and Frank, 2018). Also, HPC->PFC low gamma signals recorded during the visits to familiar objects correlated strongly with memory performances, confirming that both male and female mice with robust HPC->PFC gamma connectivity exhibit superior memory performances (Alemany-González et al., 2020). In accord, EE increases functional connectivity between the HPC and the PFC in rats and increases behaviorally dependent prefrontal rhythmicity (Ulrich et al., 2019). In summary, the neurophysiological changes identified in enriched wild-type animals combined task-independent adjustments (augmented hippocampal pyramidal activity and gamma synchrony across different brain states) and memory-dependent adjustments (enhanced theta-gamma coupling and ripples in the HPC associated with memory retrieval).

Trisomic female mice reared in impoverished environments exhibited a robust increase in theta and gamma power in the HPC and PFC across all brain states (including rest, memory performance, and sleep). In addition, a broad coupling between theta and high frequency (<80 Hz) oscillations was detected. These changes were also observed previously in TS male mice of the same transgenic line (Alemany-González et al., 2020), demonstrating similar hyper synchronicity in both genders. The excessive theta and gamma power predicted low memory performances, unraveling their pathological nature. Moreover, trisomic mice showed abnormal hippocampal-prefrontal coordination. This included reduced gamma coherence during rest, increased gamma coherence during NREM sleep and disordered directionality of signals especifically during memory retrieval. Increased HPC gamma oscillations and circuit coherence have been reported for other three partially trisomic mouse models of DS (Chang et al., 2020). Increased pyramidal activity and gamma synchrony in the HPC were prevented by EE in trisomic mice. However, theta and gamma hypersynchrony in the PFC were reduced during resting and REM sleep but not during memory acquisition/retrieval and NREM sleep. Remarkably, circuit communication ameliorated in enriched DS animals. These results put forward hippocampal hypersynchrony and hippocampal-prefrontal miscommunication as major neural mechanisms underlying the beneficial effects of EE for intellectual disability in DS.

Therefore, EE enhanced gamma synchrony in diploid mice and reduced excessive gamma in trisomic mice. How can this be reconciled? One possibility is that the hypersynchrony observed in DS animals was well above the levels of those observed in enriched diploid animals. This implies an inverted-U shaped range of gamma synchrony for optimal memory performance, whereby insufficient or excessive gamma is detrimental to memory. Another plausible explanation is that distinct neural mechanisms underlie “healthy” and “pathological” gamma synchronization. A widely accepted hypothesis is that elevation in the ratio of cellular excitation to inhibition (E-I balance), for example through increased activity in excitatory neurons or reduction in inhibitory neuron function, could give rise to increased gamma oscillations that would result in social and cognitive deficits, such as the ones observed in patients with autism and schizophrenia (Sohal and Rubenstein, 2019; Yizhar et al., 2011). Along these lines, recent findings have demonstrated that enhanced broadband gamma power can arise from asynchronous activities that result from deficient inhibition of parvalbumin-expressing neurons and increased, but desynchronized, pyramidal activity (Guyon et al., 2021). The “healthy” gamma would arise from a true synchronization of neural activity whereas the “pathological” gamma would arise from asynchronous hyperactivity. In support of this hypothesis, the spiking of excitatory neurons and LFP entropy, a measure of asynchrony, in the HPC and PFC were increased in trisomic mice. These results are in accord with the proposed hypothesis that regions of the cortex/hippocampus with reduced interneuron function and excitatory circuit compensation could be susceptible to hyperexcitability, which could lead to a pathological switch in circuit function and behavior (Sohal and Rubenstein, 2019).

In a previous study we investigated anomalous hippocampal-prefrontal neural dynamics relevant for memory processing in trisomic males and assessed the rescuing abilities of a green tea extract containing epigallocathechin-3-gallate (EGCG) (Alemany-González et al., 2020), a DYRK1A kinase inhibitor with pro-cognitive effects in individuals with DS (de la Torre et al., 2016). Here, the experiments were conducted in trisomic females, providing unique neurophysiological biomarkers of memory impairment in DS mice of both genders and the comparative rescuing effects of EE and EGCG. Trisomic females showed excessive theta and gamma synchrony in the HPC and PFC as we observed in trisomic males, thus confirming a hippocampal-prefrontal hypersynchrony in DS mice of both genders (i.e., a pathological brain state alteration). Likewise, ripple features during sleep appeared relevant for memory processing, particularly ripple power. It was amplified in diploid enriched mice but also increased in trisomic animals of both genders and rescued by EE and EGCG. It is plausible that, like the increase in gamma synchrony, enhanced and pathological ripples were generated by distinct neural mechanisms in diploid and trisomic animals. Ripples arise from the excitatory recurrent system of the CA3 region and travel to CA1, and their pattern of spiking activity is coordinated by a consortium of interneurons (Buzsáki, 2015). Pathological ripples may emerge from unbalanced inhibition of neural networks that cause broad disruptions of sleep patterns in individuals with DS and in DS mouse models (Raveau et al., 2018; Ruiz-Mejias, 2019). Finally, we identified a key biomarker that predicts recognition memory performance: HPC -> PFC low gamma directionality during memory retrieval exhibited a strong positive correlation with discrimination indices in diploid animals of both genders, was disrupted in both trisomic female and male mice, and rescued by EE and EGCG. Green tea extracts, EE and their combination restored more than 70% of the phosphoprotein deregulation in Ts65Dn mice while re-establishment of a proper epigenetic state and rescue of the kinome deregulation may contribute to the cognitive rescue induced by green tea extracts (De Toma et al., 2020). Future studies should investigate the neural substrates of a combination of EE and EGCG in diploid and trisomic animals.

In closing, both brain state adjustments and memory-associated adjustments are good candidates to underlie the beneficial effects of EE on cognition in diploid female mice. In contrast, EE prevented hippocampal and prefrontal hypersynchrony in trisomic females, suggesting distinct neural mechanisms for the generation and rescue of healthy and pathological brain synchrony, respectively, by EE.

## ACKNOWLEDGEMENTS

We are grateful to Mara Dierssen for providing scientific insight into this study and Manuel Valero for providing scripts to analyze ripple activity and entropy. This work was supported by grant 1419 from the Jerome Lejeune Foundation.

## METHODS

### Animals

Ts65Dn female mice (n = 13) and their wild-type littermates (n = 12) were obtained by breeding B6EiC4Sn.BLiA-Ts(1716)65Dn/DnJ females with C57BL/6 × 6JOlaHsd (B6C3F1/OlaHsd) hybrid males.

The parental generation was obtained from the Jackson Laboratory (Bar Harbor, ME) and a colony was generated and maintained at the Barcelona Biomedical Research Park (PRBB) Animal Facility. Mice were genotyped and ∼25% of the offspring showed trisomy. Aged-matched diploid littermates served as controls. Animals were 2 to 3 months old at the start of all experiments and weighed between 20 to 30 g. All procedures had authorization from the PRBB Animal Research Ethics Committee and the local government (Generalitat de Catalunya) and were carried out in accordance with the guidelines of the European Union Council (EU Directive 2010/63/EU) and Spanish regulations (BOE 252/34367-91, 2005).

### Environmental enrichment protocol

Female mice were housed in standard or enriched conditions for 7 weeks after weaning. The non-enriched environment consisted in standard Plexiglas cages (37x16.5x13.5 cm) where mice were housed in groups of 2 to 3, while mice in enriched conditions were housed in groups of 4 to 7 in large two-level cages (55x80x50 cm) with plastic toys that were changed every three days to maintain environmental stimulation. During the first week after completing the 7-week rearing period, mice of both groups underwent surgery for chronic electrode implantation, which was followed by one week of post-surgical recovery. All the mice were housed individually from the day of the surgery until the end of the experiment to prevent the implants from being damaged by other cagemates. Mice in the enriched group had toys in their cages that were replaced every three days until the end of the experiment.

### Surgeries

Mice were anesthetized with a mixture of ketamine/xylazine (ketamine: Imalgene 1000, Distrivet SA; xylazine: X1251-1G, Sigma-Aldrich) and placed in a stereotaxic apparatus. Anesthesia was maintained with continuous 0.5-4% isoflurane (Zoetis Spain, S.L). Small craniotomies were drilled above the medial PFC and the HPC. Five micro-screws were screwed into the skull to stabilize the implant, and the one on top of the cerebellum was used as a general ground. Three tungsten electrodes, one stereotrode and one single electrode, were implanted in the PFC and three more were implanted in the HPC. The electrodes were positioned stereotaxically in the prelimbic PFC (AP: 1.5, 2.1 mm; ML: ± 0.6, 0.25 mm; DV: -1.7 mm from bregma) and in the CA1 area of the HPC (AP: -1.8, -2.5 mm; ML: -1.3, -2.3 mm; DV: -1.15, -1.25 mm). In addition, three reference electrodes were implanted in corpus callosum and lateral ventricles (AP: 1, 0.2, -1; ML: 1, 0.8, 1.7; DV: -1.25, -1.4, -1.5, respectively). The electrodes were made by twisting a strand of tungsten wire 25 μm wide (Advent, UK), had impedances that ranged from 100 to 400 kOhm at the time of implantation and were implanted unilaterally with dental cement. Electrode wires were pinned to an adaptor to facilitate their connection to the recording system. After surgery, animals were allowed to recover during at least one week in which they were extensively monitored and received both analgesia and anti-inflammatory treatments. Additionally, animals were handled and familiarized with the implant connected to the recording cable. After the experiments ended, a mild electrical current (100 Hz, 0.1 mA, 2 s) was applied to the electrodes to mark the placement of electrode tips, which were later confirmed histologically by staining the brain slices with Cresyl violet (Sigma-Aldrich). Electrodes with tips outside the targeted areas were discarded from data analyses.

### Behavioral and Neurophysiological Characterization

We recorded single-unit activity (SUA) and local field potentials (LFPs) in freely moving mice with the multi-channel Open Ephys system at a sampling rate of 30 kHz with Intan RHD2132 amplifiers equipped with an accelerometer. We used the accelerometeŕs signals in the X, Y and Z axis to monitor general mobility of mice. We found that the variance of the instantaneous acceleration module (Acc), that quantifies the variation of movement across the three spatial dimensions, was maximal during exploration and decreased as the animals were in quiet alertness and natural sleep (Alemany-González et al., 2020; Gener et al., 2019). More specifically, raw X, Y and Z signals were first downsampled to 1 kHz, then we calculated the instantaneous module (root square[X^2^, Y^2^, Z^2^]) from which we measured the variance. In addition, recorded signals from each electrode were filtered offline to extract SUA and LFPs. SUA was estimated by first subtracting the raw signal from each electrode with the signal from a nearby referencing electrode to remove artifacts related to the animal’s movement. Next, referenced signals were filtered between 450-6000 Hz and the spikes from individual neurons were sorted using principal component analysis with Offline Sorter v4 (Plexon Inc.). Neurons with low firing rates (<3 spikes/s) were not used in the analyses. To obtain LFPs, signals were downsampled to 1 kHz, detrended and notch-filtered to remove noise line artifacts (50 Hz and its harmonics) with custom-written scripts in Python. Signals were then imported into MATLAB (MathWorks, Natick, MA). The frequency bands considered for the band-specific analyses included delta (1-4 Hz), theta (8-12 Hz), beta (18-25 Hz), low gamma (30-50 Hz), and high gamma (50-80 Hz).

### Quiet Wakefulness, NREM and REM Sleep

Recordings during quiet wakefulness were performed in an empty standard cage for 30 minutes. Briefly, low mobility was determined by a defined threshold in the output of the accelerometer signals and normal movement was defined as above that threshold. The animals typically rested for brief periods between 2 to 10 consecutive seconds. Recordings during natural sleep were implemented following the familiarization phase of the NOR task to better capture neural signals related to memory consolidation (Fig. 1a). Accelerometer measures and LFP signals from the PFC and the HPC were used to score epochs of REM and NREM sleep. NREM sleep was behaviorally defined as presenting immobility (low variations of the accelerometer) and large amplitude slow oscillations (1-4 Hz) in the PFC and HPC. Only periods of deep NREM sleep prior to REM episodes were considered for analysis. REM epochs were defined as immobility and prominent hippocampal theta rhythms. Analyses of LFPs signals during quiet wakefulness and natural sleep were performed averaging neural signals over discrete epochs of different duration (one continuous epoch per experiment; quiet wakefulness: 3 min; NREM sleep: 1 min; REM sleep: 10 s) that were chosen based on the stability of the brain state (McShane et al., 2012).

### Novel Object Recognition Test

We tested recognition memory using a well-established task that relies on mice’ innate instinct to explore novel objects in the environment (McShane et al., 2012). We used a custom-designed T-maze made of aluminum with wider and higher arms than the standard mazes (8 cm wide x 30 cm long x 20 high). The maze was shielded and grounded for electrophysiological recordings and was placed on an aluminum platform. The novel-familiar object pairs were previously validated as in (Gulinello et al., 2018). The arm of the maze where the novel object was placed was randomly chosen across experiments. The test was implemented in three phases of 10 minutes each: habituation and familiarization during the first day and long-term memory test 24 hours after familiarization. Mice were first habituated to an empty maze. Five minutes later, mice were placed again in the maze where they could explore two identical objects located at the end of the two lateral arms. Twenty-four hours later, in the test phase, mice were presented with one familiar and one novel object. Each session was videotaped via a video camera located on top of the maze. Any investigative behavior of objects, including head orientation towards the objects or sniffing at a distance below or equal to 2 cm or when the mice touched the objects with the nose, was considered object exploration. Exploratory events were identified online by looking at the video using a custom-designed joystick with a right and left button that were pressed continuously during the time of explorations. Button presses were automatically aligned to the electrophysiological file by sending TTL pulses to the acquisition system so that two more event channels were added to the recording files. Object recognition memory was defined by the discrimination index (DI) for the novel object using the difference in exploration time for the familiar object divided by the total amount of exploration of both objects (DI = [time of novel object exploration-time of familiar object exploration]/total exploration time). DIs vary between +1 and -1, where a positive score indicates more time spent with the novel object, a negative score indicates more time spent with the familiar object, and a zero score indicates a null preference (Leger et al., 2013).

LFP measures associated with memory acquisition and retrieval were obtained by averaging one-second non-overlapping windows triggered by the button presses on the joystick. Interactions with artifacts were discarded in the analyses. Each one-second window was only considered if the mouse explored the object over 600 ms. This allowed us to include many interactions from trisomic mice that were particularly short while also granting a good estimation of power at low frequencies. Memory retrieval was investigated in the 24-hour memory test during the visits to familiar objects. Memory acquisition and novelty seeking, which cannot be disentangled in this task, were investigated in the 24-hour memory test during the visits to novel objects. We found that these criteria provided the most robust results.

### Data Analyses

#### Power Spectral Analyses

We used the multitaper fast Fourier transform method (time frequency bandwidth; TW = 5 and K = 9 tapers; 1-100 Hz range; non-overlapping sliding window of 5 seconds for power spectra, 2 seconds for spectrograms, 1 second for object explorations) with the Chronux toolbox in MATLAB (Bokil et al., 2010).

#### Phase-Amplitude Modulation index

To quantify the intensity of phase-amplitude coupling we used the modulation index (MI) as described in (Tort et al., 2010). First, low frequency (delta and theta) phases were divided into 18° bins and gamma (low, high gamma and high frequency oscillations-HFO) amplitude was calculated for each phase bin. MI measures the divergence of the phase-amplitude distribution and is higher as it is further away from the uniform distribution. In order to choose specific frequency bands pairs for the MI quantification we represented overall MI in a two-dimensional pseudocolor comodulation map. A warmer color indicates coupling between the phase of the low frequency band (x axis) and the amplitude of the high frequency band (y axis) while blue depicts absence of coupling.

#### Weighted Phase Lag Index (wPLI)

To assess functional connectivity between LFPs in the PFC and HPC we used the wPLI as in (Hardmeier et al., 2014; Vinck et al., 2011), a measure of phase synchronization. First, instantaneous phases from LFP signals recorded simultaneously in both areas were determined by Hilbert transformation. wPLI measures the asymmetry in the distribution of phase differences for each frequency band between the two time-series resulting in values ranging between 0 and 1, being a higher value and a high asymmetric distribution as a consequence of a consistent phase-lag between signals in the two areas. wPLI reduces the probability of detecting false positive connectivity in the case of volume-conducted noise sources with near zero phase lag and shows higher sensitivity in detecting real phase synchronization.

#### Phase Slope Index (PSI)

To estimate the flow of information between neural signals of the PFC and HPC, we used the Phase Slope Index as in (Nolte et al., 2008) with the data2psi.m function. PSI is a robust measure based on the conceptual temporal argument supporting that the driver is earlier than the recipient and contains information about the future of the recipient. It quantifies the consistency of the direction of the change in the phase difference across frequencies. Positive slope reflects an HPC-to-PFC flow of information in a specific frequency range while a negative slope reflects the opposite.

#### Analyses of ripples during NREM sleep

Raw signals were down-sampled to 1.25 kHz and bandpass filtered between 100-600 Hz. Sharp-wave ripple events were detected by thresholding (>3 SDs) the filtered signals using ripple detection scripts from Valero et al (Valero et al., 2017). Time-frequency analysis of ripples was calculated using multitaper spectral estimation and frequency resolution of 10 Hz in the 100- 600 Hz range.

#### Analyses of entropy

To quantify the level of frequency dispersion (and consequently the level of synchronization in the neuronal activity), we performed analysis of the spectral entropy (based on Shannon entropy) of the LFP (30-150 Hz), where a narrower power distribution results in lower entropy values, and a large power distribution results in higher entropy values, with no dependency on the LFP frequency (Valero et al., 2017).

#### Statistical Analyses

All data is represented as the mean ± SEM. We used unpaired T tests to compare behavioral measures between genotypes. Paired T tests and mixed ANOVAs were used to assess the effects of environmental enrichment across brain regions. Two-WAY ANOVAs were used to assess the effects of environmental enrichment across genotypes. Mann-Whitney non-parametric tests were used to assess the effects of enrichment on phase-amplitude coupling (PAC), entropy, phase coherence (wPLI) and directionality of signals (PSI) between brain regions. We used Bonferroni correction or Tukey HSD post-hoc tests in the ANOVAs when appropriate. To identify significant correlations between neurophysiological measures and DIs, Pearson or Spearman correlations were used for parametric and non-parametric distributions, respectively. Statistical analyses were implemented with the statistical package SPSS.

